# Electrical synapse asymmetry results from, and masks, neuronal heterogeneity

**DOI:** 10.1101/2021.06.30.450525

**Authors:** Austin Mendoza, Julie S. Haas

## Abstract

Electrical synapses couple inhibitory neurons across the brain, underlying a variety of functions that are modifiable by activity. Despite recent advances, many basic functions and contributions of electrical synapses within neural circuitry remain underappreciated. Among these is the source and impact of electrical synapse asymmetry. Using multi-compartmental models of neurons coupled through dendritic electrical synapses, we investigated intrinsic factors that contribute to synaptic asymmetry and that result in modulation of spike time between coupled cells. We show that electrical synapse location along a dendrite, input resistance, internal dendritic resistance, or directional conduction of the electrical synapse itself each alter asymmetry as measured by coupling between cell somas. Conversely, true synapse asymmetry can be masked by each of these properties. Furthermore, we show that asymmetry alters the spiking timing and latency of coupled cells by up to tens of milliseconds, depending on direction of conduction or dendritic location of the electrical synapse. These simulations illustrate that causes of asymmetry are multifactorial, may not be apparent in somatic measurements of electrical coupling, influence dendritic processing, and produce a variety of outcomes on spike timing of coupled cells. Our findings highlight aspects of electrical synapses that should be considered in experimental demonstrations of coupling, and when assembling networks containing electrical synapses.

## INTRODUCTION

Electrical synapses represent a major form of communication between GABAergic neurons across neuronal tissue, with many basic properties that have not been extensively explored. Rectification, or asymmetry of transmission, is a frequently noted aspect of electrical synapses: it is the property of unequal transmission of electrical signals between two neurons. Electrical synapses have been well-studied in invertebrates, where evidence of asymmetry comes from species including crayfish (Furshpan & Potter, 1959), drosophila giant fibers (Phelan et al., 2008), lobster stomatogastric nucleus (Johnson et al., 1993), and the C. elegans escape circuit (Liu et al., 2017; Shui et al., 2020). In invertebrate systems asymmetry varies widely, with some synapses displaying full rectification. In contrast, asymmetry at synapses between mammalian neurons is often smaller. Demonstrations of electrical synapse asymmetry are numerous throughout the mammalian brain, including retina (Veruki & Hartveit, 2002), cortex (Galarreta & Hestrin, 2002), inferior olive (Devor & Yarom, 2002), dorsal cochlear nucleus (Apostolides & Trussell, 2013), mesencephalic trigeminal nucleus (Curti et al., 2012), cerebellar Golgi cells (Dugué et al., 2009; Szoboszlay et al., 2016), and the thalamic reticular nucleus (TRN)(Haas et al., 2011; Sevetson & Haas, 2015; Zolnik & Connors, 2016). Recent results show that asymmetry can be modified during the activity that results in electrical synapse plasticity (Fricker et al., 2021; Haas et al., 2011), indicating that it is a dynamic property that is under activity-dependent regulation.

Asymmetry of electrical transmission can in principle result from a wide variety of influences. It has been well established in non-mammalian systems that directional differences in conductance between two coupled cells can result from heterogeneity or heterotypy of gap junction plaques or hemichannels in each membrane (Bukauskas et al., 1995; Rash et al., 2013), or from differences in hemichannel protein scaffolding (Marsh et al., 2017). While hemichannel differences have been demonstrated in HeLa cells expressing connexin isoforms (Bukauskas et al., 1995) and at the goldfish mixed synapse onto Mauthner cells (Rash et al., 2013). Connexin-sourced asymmetry is thought to be unlikely for neuronal mammal synapses, as connexin36 does not oligomerize with other connexins (Li et al., 2004; Teubner et al., 2013), and in expression systems appears to form perfectly symmetric synapses (Srinivas et al., 1999). However, residual coupling has been noted between TRN neurons in connexin36 knockout mice, and that coupling was more asymmetrical (Zolnik & Connors, 2016). Large gradients of Mg^2+^ concentration also produce asymmetric signaling for neuronal synapses (Palacios-Prado et al., 2013). Additionally, gating properties of connexin channels can produce asymmetry in computational models (Snipas et al., 2017). Macroscopic differences in coupled neurons, such as differences in input resistance have been proposed as a straightforward reason that one would observe asymmetry in coupling coefficients (Fortier, 2010; Veruki & Hartveit, 2002), but for TRN synapses, asymmetry remains even after computing estimates of conductance that should in principle minimize contributions of input resistance (Haas et al., 2011; Sevetson & Haas, 2015). While many reports of electrical synapses across the mammalian brain include asymmetry in their measurements, some reports do not note it, or only note that in their observations, synapses were symmetrical as expected. In all, asymmetry and its sources remain underexamined at mammalian electrical synapses.

Beyond observations, the functional consequences of electrical synapse asymmetry on neural activity are not robustly understood. Electrical synapses have been widely shown to contribute towards synchrony in neuronal networks in both computational models (Chow & Kopell, 2000; Gutierrez et al., 2013; Lewis & Rinzel, 2003; Pfeuty et al., 2005; Traub et al., 2005) and experimentally (Long, 2004; Mann-Metzer & Yarom, 1999; Veruki & Hartveit, 2002; Vervaeke et al., 2010), and oscillations are more robust when electrical synapses take on asymmetric values (Gutierrez & Marder, 2013). In non-rhythmic settings, strong asymmetry can produce nearly unidirectional communication that serves to reliably excite one coupled cell, as is the case with the club endings onto Mauthner cells in goldfish (Rash et al., 2013), and dorsal cochlear nucleus (Apostolides & Trussell, 2013). Dendritic electrical synapses are proposed to interact with nearby GABAergic synapse activity through the shunting of current through Cl^-^ channels (Lang et al., 1996; Llinas et al., 1974). Additionally, in C. elegans coupled motor neurons, electrical synapses spread excitation during contraction and inhibit cell pairs between cycles through a shunting effect (Choi et al., 2021). Electrical synapses modulate spike times in coupled neighbors in TRN by up to tens of milliseconds (Haas, 2015; Sevetson & Haas, 2015), and asymmetric coupling can add to that modulation, even reversing firing order between two coupled cells receiving closely-timed inputs (Sevetson & Haas, 2015). In a model thalamocortical circuit, coupling amongst feedback inhibitory neurons enhances discrimination of inputs sent to cortex by relay cells (Pham & Haas, 2018). In a canonical model circuit with feedforward inhibition, electrical synapses enhance subthreshold integration (Pham & Haas, 2019). In a toadfish vocal circuit, electrical coupling between feedforward inhibitory neurons enhances synchrony and temporal precision (Chagnaud et al., 2021) and a similar effect occurs for cerebellar basket cells (Hoehne et al., 2020). However, few models have yet included electrical synapses in morphologically extended cells, let alone asymmetrical synapses.

Here we used compartmental models of coupled TRN neurons to investigate how fundamental neuronal properties contribute to electrical synapse asymmetry, including synapse location, strength, direction of conductance, dendritic geometry, and input resistance. We show that a variety of these factors can produce apparent differences in coupling coefficients as measured between somas. Together, these results underline that one should in fact always expect asymmetry at almost any electrical synapse. Conversely, these same properties can mask true asymmetries of electrical conductance, resulting in apparently equal coupling as measured between somas. Further, we show that asymmetry is a powerful regulator of spike timing in coupled neurons, and electrical synapses between dendrites can exert locally powerful influence that is not readily apparent at the soma, highlighting the necessity of including electrical synapses in morphologically detailed models, circuits or connectomes.

## RESULTS

To address the impact of neuronal excitability and morphology on electrical synapse communication, we built a three-compartment TRN cell model based on those previously used (McCormick & Huguenard, 1992; Pham & Haas, 2018, 2019; Traub et al., 2005). To validate the model’s dendritic geometry, we used coupling between compartments that generated reasonable amplitudes of backpropagated single spikes (Figure 1A), and sublinear dendritic responses to AMPAergic current injections (Figure 1B) (Connelly et al., 2017). We then added electrical synapses between matched compartments of two identical TRN model cells and measured the coupling coefficients resulting from hyperpolarizing current applied to and measured at the somas (Fig. 1C_1_). For matched-compartment connections, electrical synapses produced higher coupling coefficients when synapses were closer to the soma, as the current between electrodes has a more direct path (Figure 1D). A single-compartmental model used for comparison (Fig. 1C-D, black) resulted in stronger coupling coefficients, as introducing dendrites imposed a load upon the circuit. Similarly, when current is applied and coupling measurements are taken between distal dendrites (Figure 1C_2_), coupling is stronger for dendritically located synapses but decreases as the synapse location approaches the soma (Figure 1D_2_). This is a simple result of cable properties, but highlights the notion that electrical synapses may produce strong and effective coupling between dendritic compartments that is not apparent from somatic measurements.

**Figure 1.**
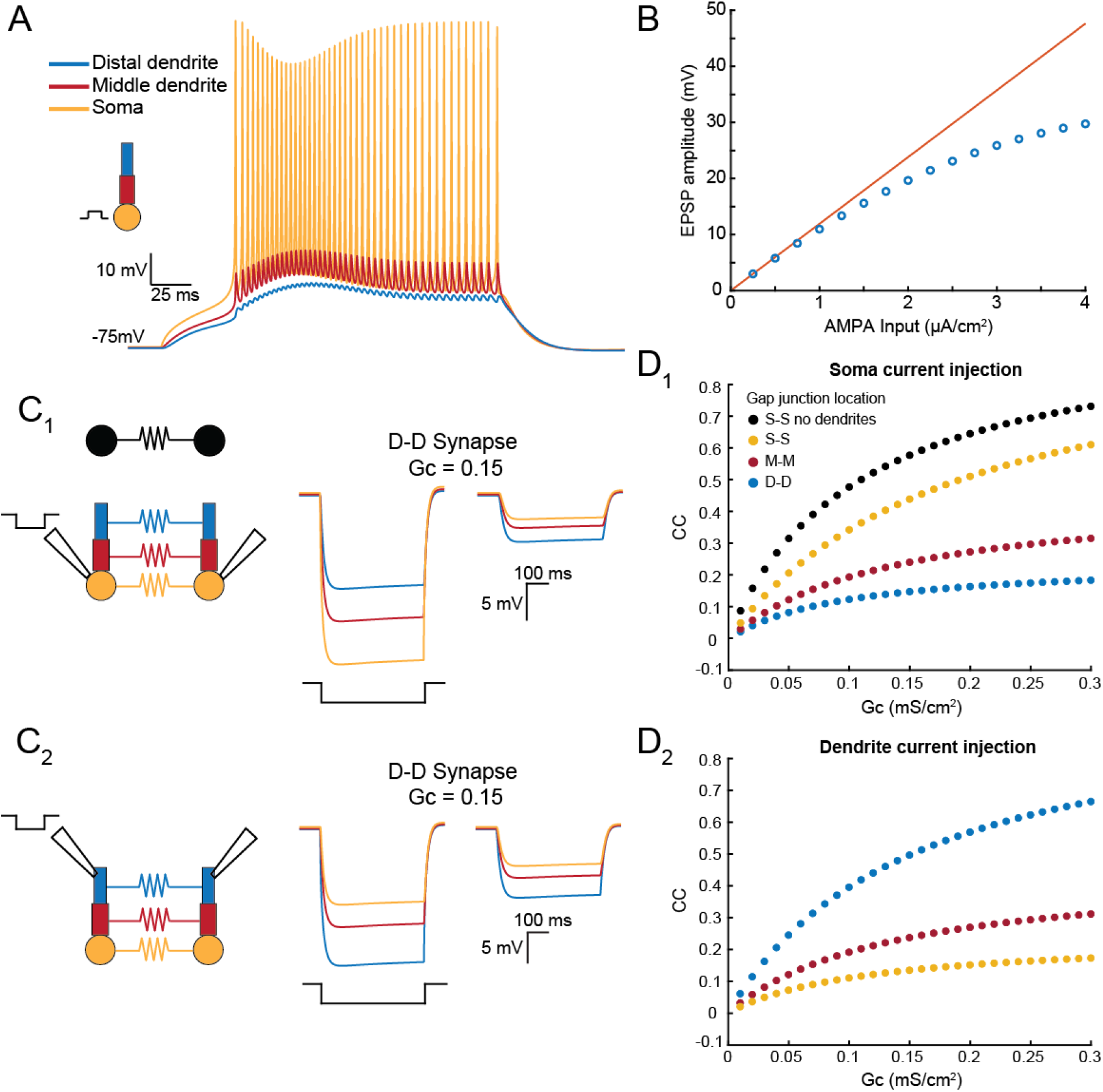
Compartmental TRN model and characterization of coupling responses for gap junctions of varied dendritic locations. A) Schematic of compartmental TRN model and representative traces from each compartment in response to square current injection at the soma compartment. B) Postsynaptic response to an AMPAergic input delivered to distal dendrite. EPSP amplitudes were sublinear above 2 µA, due to lack of active conductances in dendrites. C) Schematic for electrically coupled models. Voltage traces in both cells result from a hyperpolarizing current injections into the soma of cell 1 (top traces) or injection into dendrites (bottom traces). D) Coupling coefficients (cc) measured as in (C) for GJs located between the somas (yellow), middle compartments (red), and distal dendrites (blue). An identical single-compartment model is shown for comparison (black).

Next, we varied the location of electrical synapses between the dendritic compartments of each cell, and again measured coupling between somas of cell 1 and cell 2. In all cases, the coupling for mixed-location synapses was intermediate to the values obtained for connections between matched compartments (Figure 2A-C). Interestingly, coupling values for pairs of compartment connections that one might initially expect to produce the same coupling, such as soma-middle (S-M) and middle-soma (M-S) (Fig. 2A, orange dots), do not produce identical coupling coefficients. The source of asymmetry in this case is the differences in dendritic load that siphon soma-applied current from the gap junction pathway. Specifically, for S-M connections, the dendritic load to soma-applied current comprises resistance from both M and D compartments and is larger than the remaining dendritic load from a single D compartment when current is applied to the opposite soma. These differences lead to differences in the currents crossing the gap junction when current is separately applied to each soma, and thus the coupling coefficients are asymmetric as measured between somas. Comparing Fig. 2A-c, we note that this effect is strongest for connections closer to the soma, where the differences in dendrites distal to the electrical synapse are largest.

**Figure 2.**
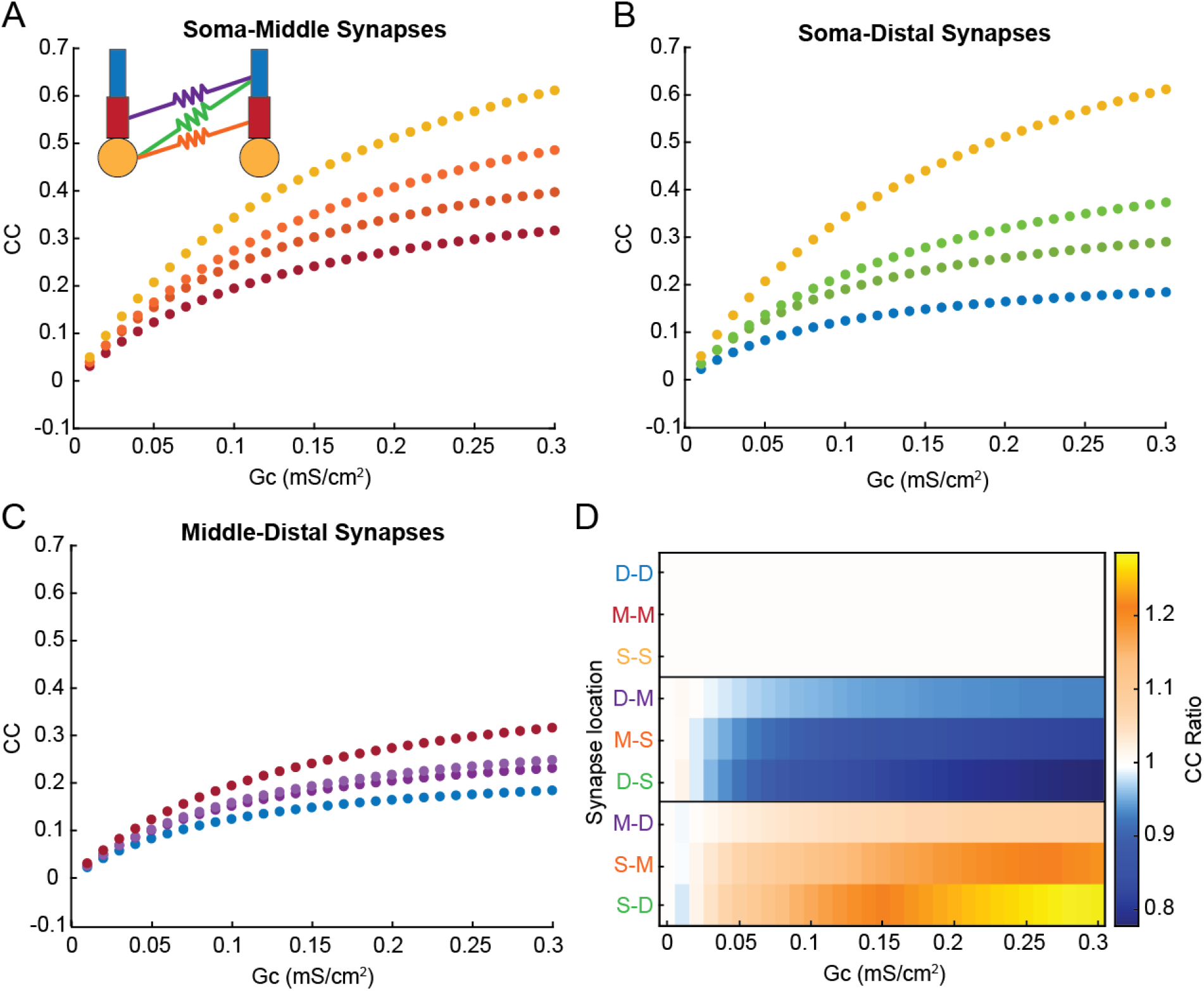
Electrical synapse location and strength contribute to coupling coefficient and asymmetry measured at the soma. A) Coupling coefficients measured from cell 1 to cell 2 for all sets of somatic and middle-compartment electrical synapses. Soma-soma synapses are in yellow, middle-middle synapses are in red, and both types of soma-middle synapses are orange.. B) as in (A) for somatic and distal synapses. C) as in (A) for middle and distal synapses; note overall decrease in coupling for more-distant synapses. D) Coupling coefficient ratio (cc_12_/cc_21_) for each synapse location and strength. Locations are grouped by effect; the top box shows GJs between matched compartments that are symmetric, as expected. In the middle box, the GJ was closer to the soma of cell 2, and thus cc_21_ was larger. In the bottom box, the GJ was closer to the soma of cell 1, and thus cc_12_ was larger. Asymmetry increased with difference between location of the GJ, and with proximity to the soma.

We compared asymmetry, or cc ratios, for all synapse locations as a function of synapse strength (Figure 2D). As expected, connections between the same compartments (eg M-M) were perfectly symmetrical for all values of electrical synapse conductance. In contrast, mismatched synapses are marked by decreasing or increasing cc ratios, with the mirror cases producing the same degree of asymmetry in opposite directions (eg M-D and D-M). We also noted that asymmetry was greater for more mismatched synapse pairs (eg S-D and M-D, or the blocks in Fig. 2D), as the distal-soma connections produce cc ratios furthest from 1. Further, asymmetry was greatest for synapses connected to the soma, while middle-distal synapses showed lower values of asymmetry. These simulations demonstrate that apparent or effective asymmetry between somatic integrators can arise from difference in synapse location, when perfectly symmetrical electrical synapses encounter asymmetrical spatial differences between identical somas and dendrites, and thereby dictate effective asymmetry.

Effective asymmetry could also arise from differences in basic excitability, e.g., membrane input resistance R_in_. We altered R_in_ by changing leak conductance in cell 2 of the model (Fig. 3A), and measuring coupling coefficient cc in both directions. When GJs coupled two somas of differing R_in_, cc was determined only by R_in_ of cell 2 (Fig. 3B, yellow); cc_12_ varied, while cc_21_ stayed constant. As GJs were more distant from the soma, voltage divisions allowed both cc_12_ and cc_21_ to change, though changes in cc_12_ were always larger. Differences in GJ location also contributed to asymmetry here, again splitting the differences between the extremes, similarly to the effect shown in Fig. 2.

**Figure 3.**
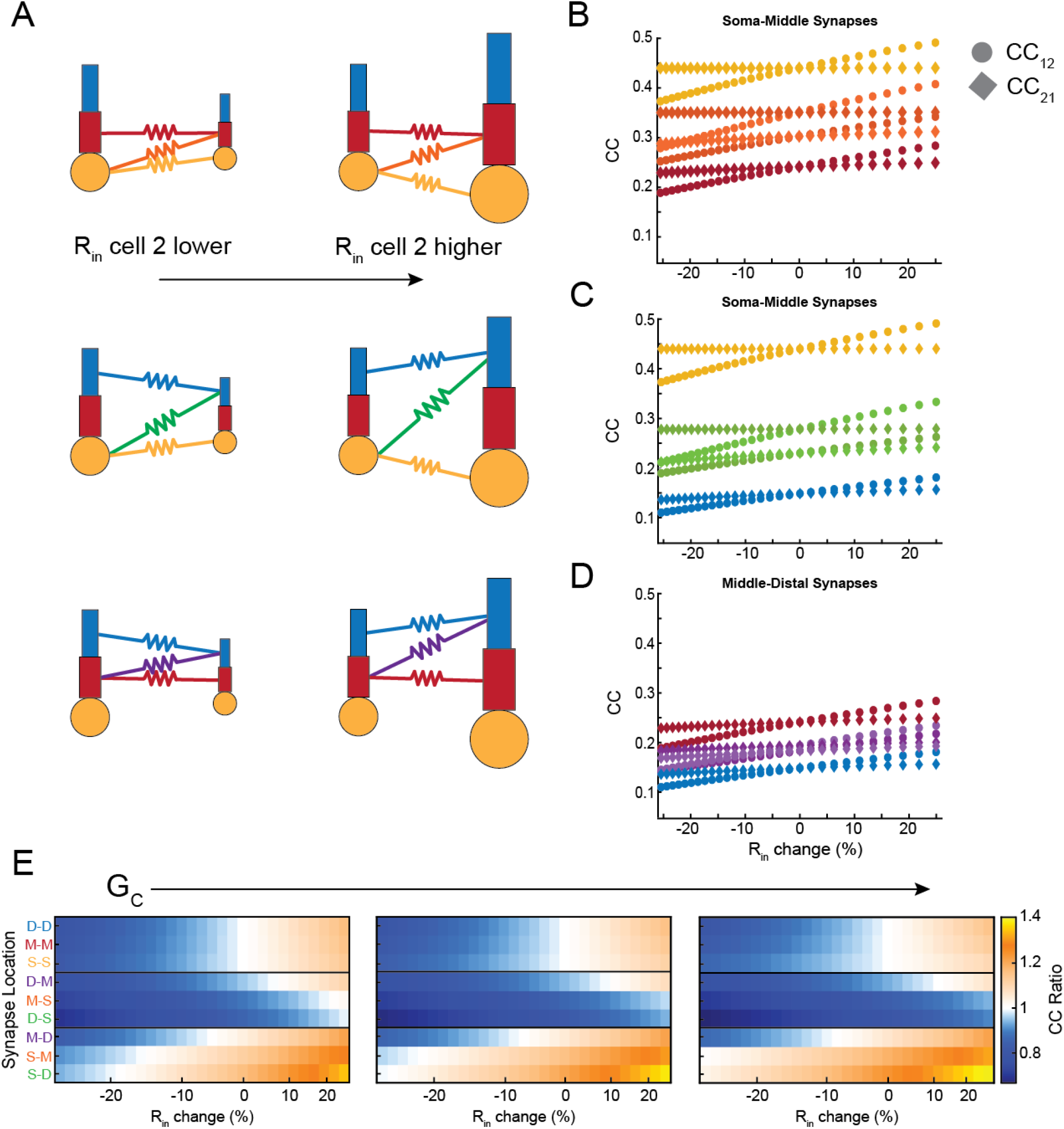
Dependence of asymmetry on synapse location and input resistance differences. A) Schematics depicting differences in input resistance, and varied synapse locations. B) Coupling versus difference in input resistance for GJs (G_C_ = 0.15 mS/cm^2^) between distal and middle compartments. Difference in input electrical synapses resistance are expressed as cell 2 relative to cell 1. C) As in (B) for GJs between middle-middle, middle-soma and soma-soma compartments. D) As in (B) for GJs between distal-distal, distal-soma, ands soma-soma compartments. E) Asymmetry plotted against input resistance differences, grouped by electrical synapse strength G_C_, which increases across panels: 0.1, 0.15 and 0.2 mS/cm^2^.

These effects are summarized in Fig. 3E, which shows simulations for three values of electrical synapse strength G_C_ for all synapse locations and input resistance mismatches. For matched-compartment locations (top boxes), asymmetry was determined only by differences in R_in_. For GJs that coupled mismatched cells at mismatched locations, synapse location appeared to be a weaker effect than input resistance mismatch: the cell with the GJ closer to its soma always yielded a smaller coupling (e.g. middle box: synapses are closer to soma 2, and produced asymmetry < 1). As in Fig. 2, asymmetry was strongest for synapses closer to the soma. These simulations showed us that increasing G_C_ amplified the asymmetry produced by differences in R_in_. Synapses that were mostly below one in cc ratio further decreased in cc ratio (Fig. 3E, middle rows), while locations with cc ratio above one increased with the strength of the synapse (Fig. 3E, bottom rows). Synapses between similar compartments (top rows) showed minimal changes with increasing strength of the synapse.

To further examine how heterogeneity between two coupled cells could contribute to effective asymmetry, we altered the internal coupling conductance between compartments of the cell. For all synapse locations, differences in dendritic coupling altered resulting cc ratios (Figure 4), but by amounts smaller than synapse mismatch or input resistance difference. Increasing dendritic conductance favors transmission into that cell and thus lowers cc ratio when cell 2 has more conductive dendrites. Similarly, cc ratio increases when cell 1 is higher in dendritic conductance. This result is consistent for the connections between same cellular compartments, which are symmetric when morphology is the same, and the mismatched locations which are asymmetric in the same case. Though morphology may not produce substantial asymmetry alone, in conjunction with synapse location the intrinsic differences between two cells will fine-tune the overall coupling and asymmetry measured between them.

**Figure 4.**
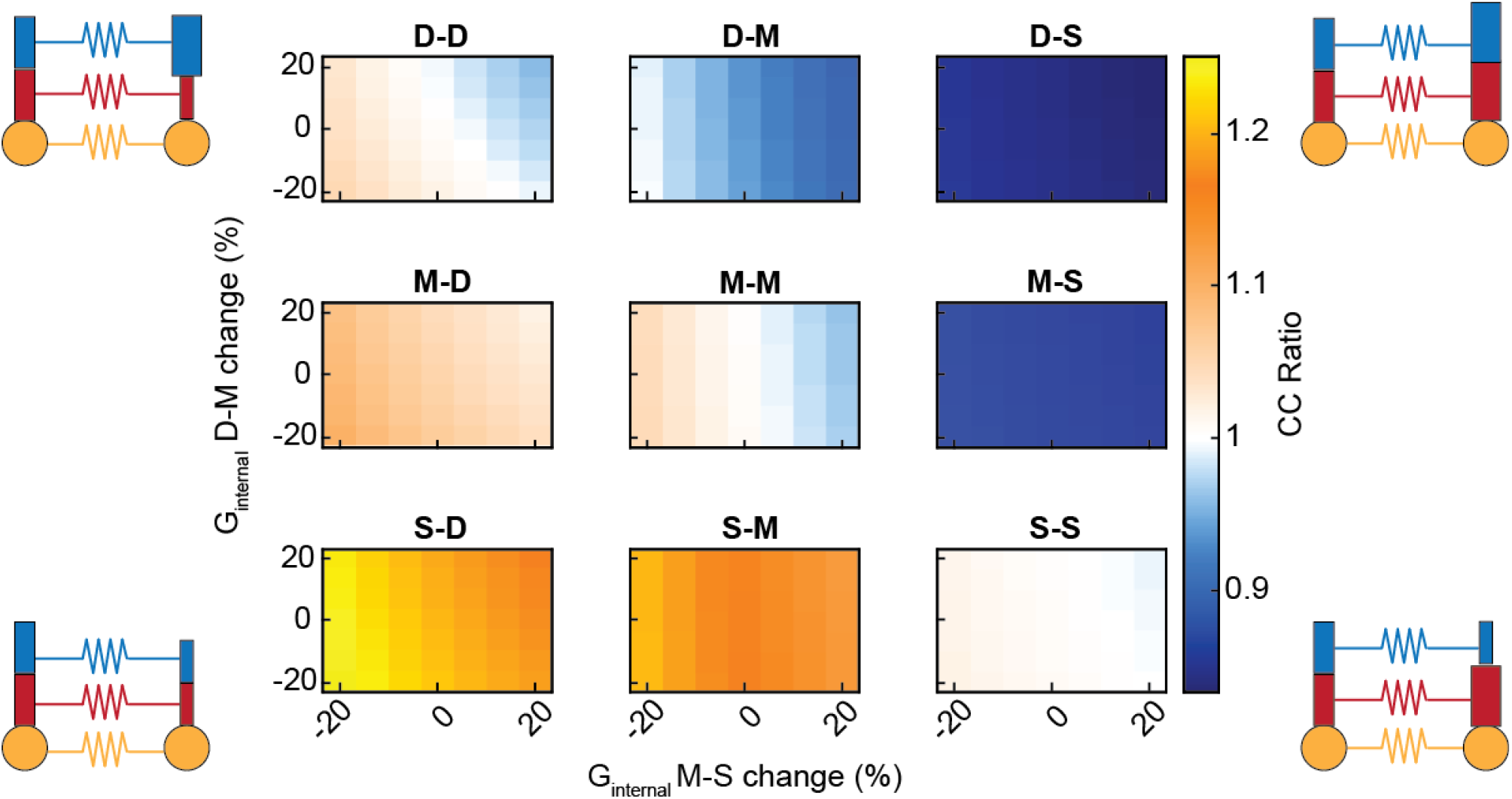
Differences in dendritic morphology fine-tune coupling coefficients and asymmetry. Internal conductances between the three subcellular compartments were altered in cell 2, as shown in schematics. Changes in middle-soma conductance is plotted on the x-axis, and change in distal-middle conductance is plotted on the y-axis. cc ratio is represented by the heat map. In all cases, decreasing internal conductance increased cc ratio, while increasing internal conductance in decreased cc ratio. Gc was 0.15 mS/cm^2^ for all synapses in these simulations.

The previous set of simulations used a symmetrical synapse to show that several aspects of cellular properties and synapse locations that can yield apparent asymmetry. Next, we asked whether an electrical synapse that was itself asymmetrical could produce the same effective or apparent asymmetries. We varied the conductance G_C_ electrical synapse between somatic compartments, and again examined the effect of input resistance changes on apparent asymmetry. Our results demonstrate that similar values of effective asymmetry could arise from either G_C_ ratio or input resistance difference (Figure 5). For one pairing of input resistance (column in Fig. 5), synaptic asymmetry could produce a range of effective asymmetry. These simulations illustrate the potential for cc values recorded from the soma to appear similarly asymmetric, whether asymmetry is produced from differential conductivity or synapses themselves.

**Figure 5.**
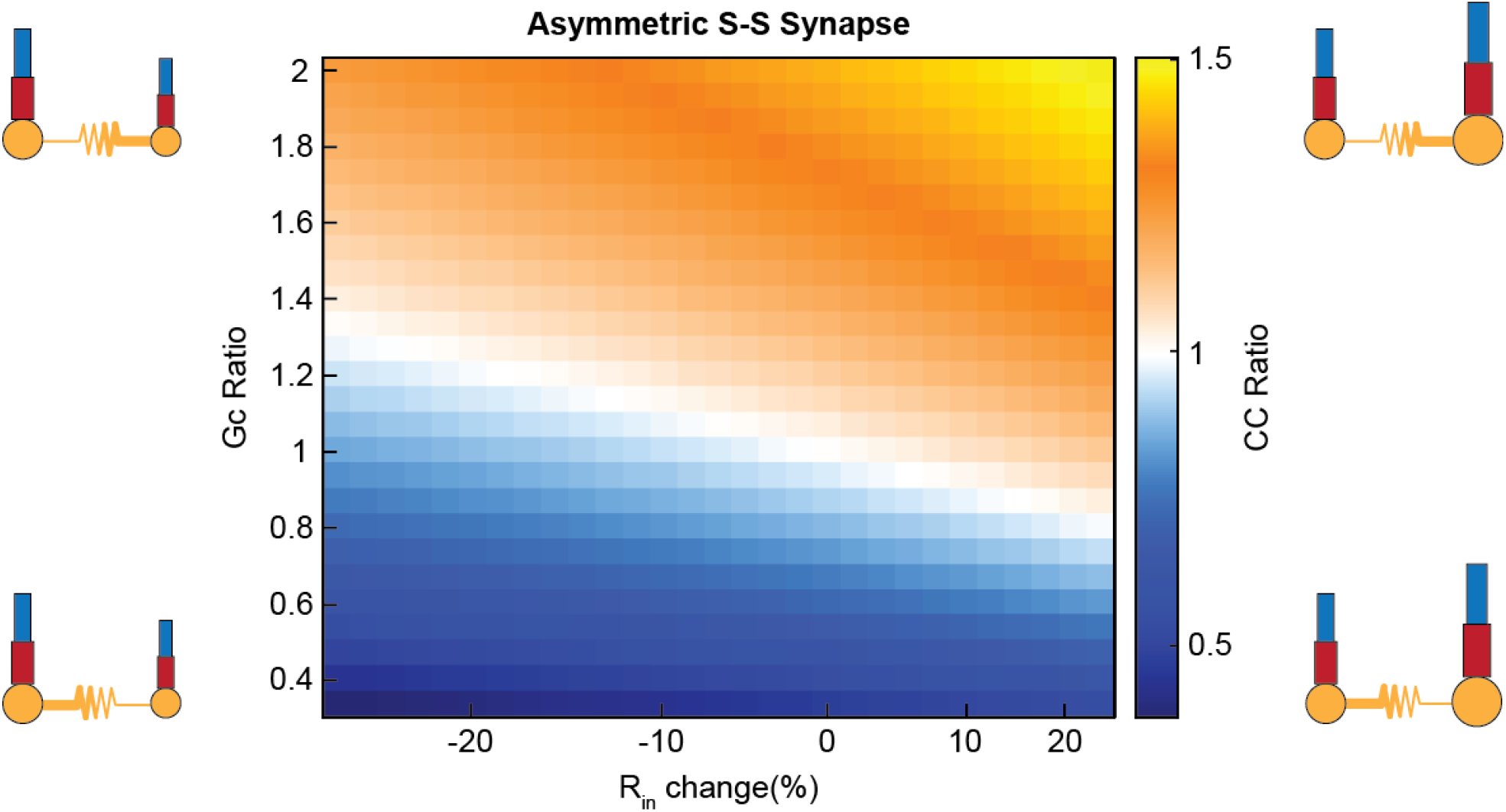
Asymmetry due to altering directional conductance produces similar5 response in coupling coefficients compared to synapse location asymmetry. Directional conductance changes between the cells produces a similar degree of asymmetry in cc ratio, with input resistance difference between the two cells predictably shifts the cc ratio values. Gc ratio (Gc_12_/Gc_21_) from altering conductance from cell 1 to cell 2 was varied while the opposite direction was held constant (Gc_21_ = 0.15 mS/cm^2^). Input resistance was altered in cell 2 relative to cell 1. Neuron schematics depict direction with larger conductance as a thicker resistor symbol, while size of the cell indicates change in input resistance.

Together, the previous results show that asymmetry in coupling as measured between somas can arise from a number of factors. We demonstrate this masking in Figure 6: the same amount of asymmetry in coupling as measured at the soma can arise from independent sources. Higher transmission to cell 2 by the same proportion (cc ratio ∼ 1.2) can be produced by asymmetric gap junction with Gc ratio of 1.8 and R_in_ change of -20% (Figure 6B), or M-D synapse location and +25% R_in_ change (Figure 6D), or S-M synapse with higher dendritic conductance in cell 1 (Figure 6F). Alternatively, higher transmission to cell 1 (cc ratio ∼ 0.8) can be produced by asymmetric gap junction with Gc ratio of 0.67 and +6% R_in_ change (Figure 6C), or M-S synapse and -12% Rin change (Figure 6E), or D-S synapse with higher dendritic conductance in cell 2 (Figure 6G). Thus, asymmetry measured at the soma is not informative as to its source, and more pertinently, fails to provide insight into processing in coupled dendrites.

**Figure 6.**
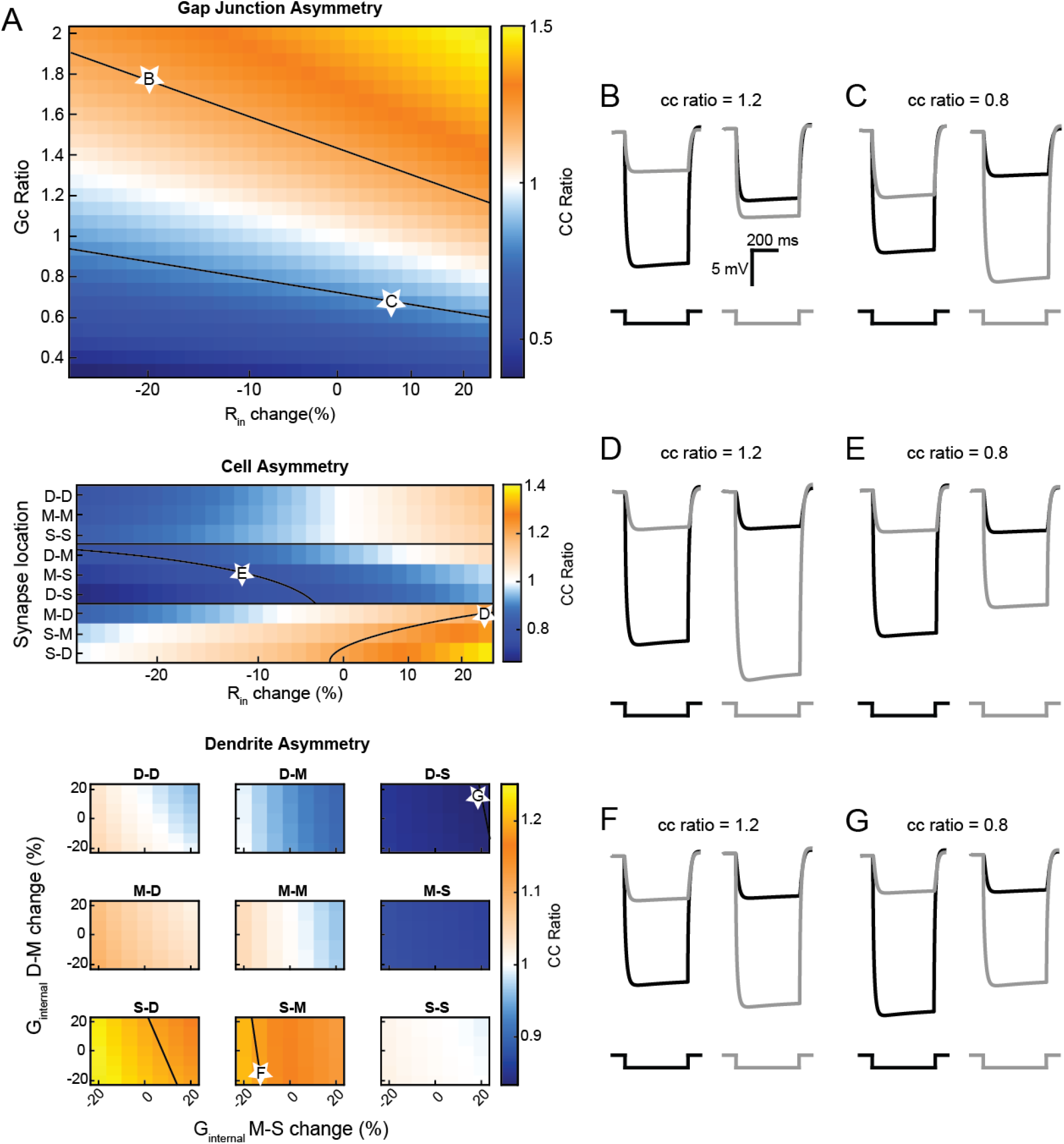
Varied scenarios can produce identical asymmetry as measured at the soma. A) Approximate isoclines (black lines) show parameters that produce the same degree of asymmetry (cc ratio of 1±0.2). Representative traces (B-G) were taken from data along these isoclines (white stars) for cases of directionally asymmetrical conducting gap junctions, differing synapse locations, or differing dendritic geometries. B) G_C_ ratio 1.8, R_in_ change - 20%. C) G_C_ ratio 0.667, R_in_ change +6%. D) M-D GJ: R_in_ change +25%. E) M-S GJ, R_in_ difference -12%. F) S-M GJ, G_MS_ change -13.3%, G_DM_ change -20%. G) D-S GJ, G_MS_ difference +20%, G_DM_ difference +20%.

Finally, we examined the impact of asymmetry on the function of coupled pairs. Electrical synapses have been previously shown to modulate latency of action potentials in coupled pairs (Haas, 2015; Sevetson & Haas, 2015). We measured latency to a burst-like input pattern of AMPAergic synaptic currents delivered to distal dendrites of a coupled pair, in order to mimic excitatory afferent activity received by cells of the thalamic reticular nucleus (Gentet & Ulrich, 2003) from bursting thalamic relay cells (Figure 7A). We tested each synapse location and delivered AMPAergic bursts to the two cell pairs with increasing difference in burst onset time between the two cells. For synapses between the similar compartments (Figure 7B, D-D), latency difference increases with G_C_ and difference in input time. Dissimilar locations alter the latency modulation with a variety of effects, with synapses in varied locations shifting latency modulation curves either up (D-M) or down (M-D), producing a variety of outcomes in cell firing by as much as 20 ms. This trend is generalized in the asymmetrically conducting synapse, where G_C_ ratio >1 produces higher latency modulation, and values <1 shift latency modulation lower (Figure 7C).

**Figure 7.**
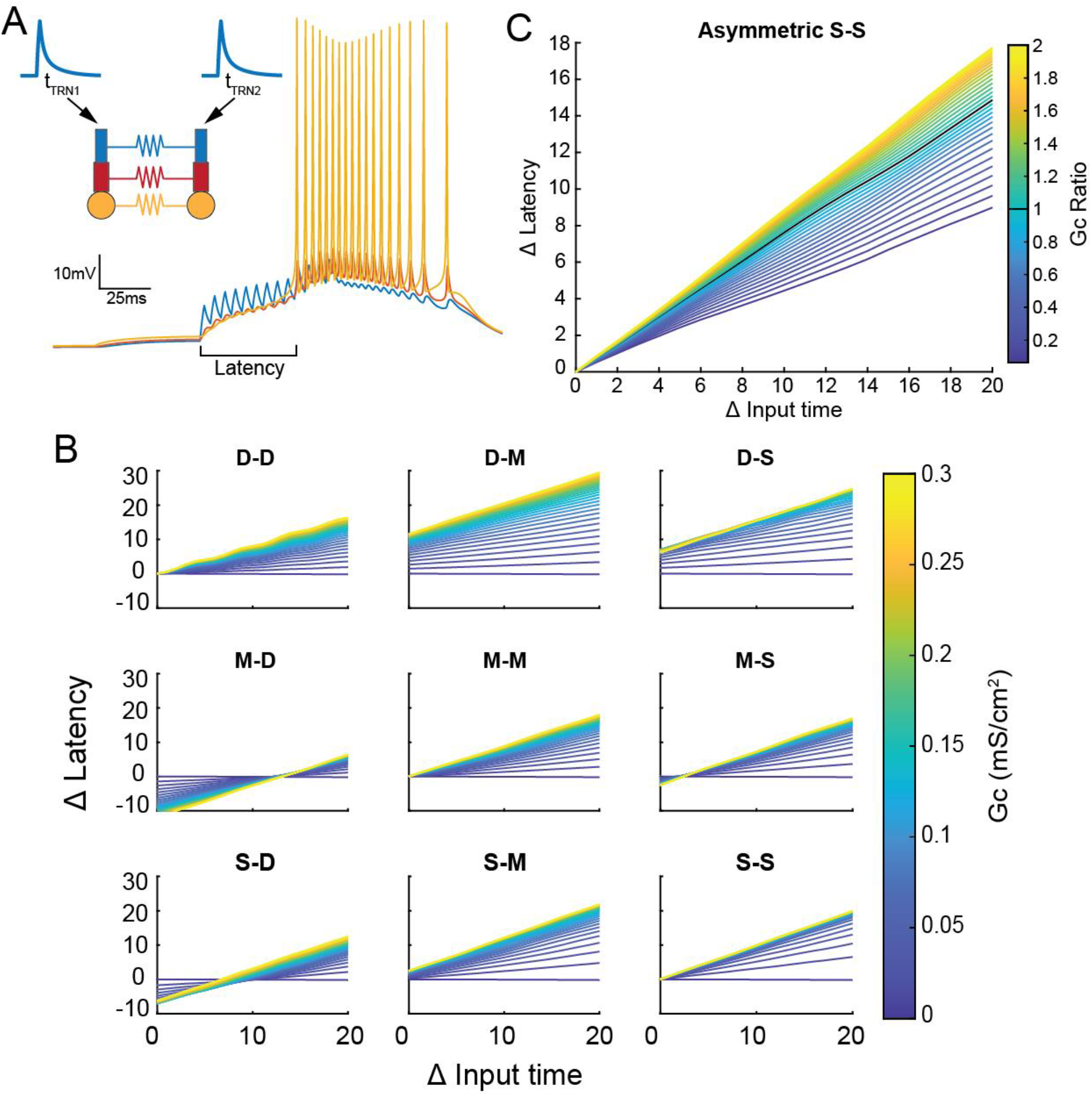
Asymmetry strongly alters the latency modulation produced by electrical synapses. A) Two electrically coupled cells received AMPAergic burst-like inputs (13 events with 1 µA amplitude, 5 ms ISI) with differing onset times. B) Changes in latency for varied electrical synapse strengths and input time differences. C) An asymmetrically conducting somatic gap junction similarly shifts latency modulation curves higher for G_C_ ratio greater than 1 and shifts down when G_C_ ratio is lower than 1.

## DISCUSSION

Asymmetry of transmission at electrical synapses has been widely noted but rarely explored in depth, perhaps due to the experimental difficulties of identifying and localizing specific GJs *in vitro* or *in vivo*. Nonetheless, because asymmetry is pervasive and can result in extreme cases in which spikes in one cell more or less faithfully drives spiking in the coupled neighbor (Apostolides & Trussell, 2013; Rash et al., 2013), we sought to understand how basic neuronal properties could influence effective coupling, and thereby the function of coupled networks. Here we have shown that asymmetry can arise from a variety of intrinsic differences in neuronal properties as well as differences in subcellular localization of the gap junction between somas. In practice, asymmetry is a combined product of all of these factors together. We also found that asymmetrical and/or strong synapses between dendritic compartments can be masked from somatic detection by the same properties. Our measurements here focused on soma-to-soma transmission, as ultimately, asymmetry between somas is the last stop before spike generation in the AIS, and because electrical synapse strength is traditionally measured between somas. Indeed, we have also shown that asymmetry substantially impacts spike times in coupled pairs.

Precise locations of electrical synapses along dendrites have proven difficult to determine. Average intersomatic distance between cells in TRN are ∼100 µm (Lee et al., 2014) suggests more distal locations on TRN dendrites are more likely positions to make close contact between cells. In coupled interneurons of cortex layer IV, synapses are located all along the dendrites, and measurements from 204 cells showed strong asymmetry in localization, with 90% synapses within 50 - 75 µm of one soma, but up to 250 µm away from the coupled soma. Asymmetrical localization also appears to be a feature of coupling between Golgi cells (Szoboszlay et al., 2016). The strongly asymmetrical synapses of the DCN also appear to couple mismatched distances from fusiform and stellate somas (Apostolides & Trussell, 2013) In brainstem MesV cells, GJs are appear to be located at or very close to the soma (Curti et al., 2012). In contrast, in inferior olive (Devor & Yarom, 2002; Hoge et al., 2011) cells are coupled at quite distal dendrites, such that somatic measurements of coupling themselves are quite small. Asymmetry has also been shown to amplify EPSPs in mixed synapses (Liu et al., 2017). Dendritic integration is likely to be influenced by the presence of gap junctions along dendrites, as morphology determines firing patterns of biophysical models (Mainen & Sejnowski, 1996), and as our results demonstrate, substantial coupling influences on dendritic processing may not be appreciably indicated by somatic measurements.

Effective asymmetry results in differentially directed signal and information flow through a network that includes realistic electrical coupling. Our results here raise interesting questions whether coupled networks actively and programmatically regulate any of the factors that result in asymmetry in order to produce precise direction of information flow within a circuit or as a function of activity state. Increasing electrical synapse strength through trafficking of connexin proteins, a process which is controlled by cAMP expression (Palumbos et al., 2021), may produce or effectively relocate a synapse slightly closer to or distal from the soma. Distances of dendritically located electrical synapses between cerebellar Golgi cells do not correlate with coupling strength measured between somas (Szoboszlay et al., 2016), indicating a possible compensation for distance by strength upregulation for those cells. Further, our previous work demonstrating activity-dependent plasticity showed that asymmetry changes systematically with unidirectional activity or ion flow across the gap junction (Fricker et al., 2021; Haas et al., 2011). Those results imply that asymmetry is a modifiable element of electrical synapse plasticity. Our results here also point out that cellular changes, such as activity-induced changes in dendritic resistance or mutation-induced localization of GJs, could result in the changes in asymmetry measured, in addition to the possibility of changing the conductance itself.

Asymmetry, as it influences spike times in coupled cells, has downstream effects on the synaptic targets of the coupled cells.. Symmetrical electrical synapses between model TRN cells act to merge TC spike times in response to inputs of similar strength or timing, or can separate spikes from dissimilar inputs (Pham & Haas, 2018). We hypothesize that TRN neurons with asymmetric gap junctions will inhibit thalamocortical relay cells unequally, shifting the balance between merging or distinguishing signals as they are relayed to cortex. Including asymmetry as a factor in TRN networks will be important to understanding how TRN cells orchestrate the attentional spotlight at sensory thalamic nuclei. In canonical feedforward circuits, coupling between inhibitory interneurons impacts integration in principal cells (Pham & Haas, 2019). Recent investigations further show the influence of electrical synapses on temporally precise inhibition in feedforward circuits (Chagnaud et al., 2021; Hoehne et al., 2020). Asymmetry, as it can be applied to electrical synapses in these general motifs may impact the many gap junction coupled feedforward and feedback circuits that embed electrical synapses across the brain.

## MATERIALS AND METHODS

### Modelling

Models were built upon those previously reported in (Pham & Haas, 2018). We use Hodgkin-Huxley formalism solved by a second order Runge-Kutta ODE solver in MATLAB (Mathworks). Model TRN cells are comprised of a three-compartment model consisting of one soma and two dendritic compartments, approximating the middle and distal regions of the dendrite. Compartments were connected by a static conductance of 0.35 between distal and middle dendrites, and 0.4 between middle dendrite and soma. Membrane capacitance was 1.2 µF/cm^2^. Currents included in our model were fast transient Na+, K+ delayed rectifier (Kd), K+ transient A (Kt), slowly inactivating K (K2), slow anomalous rectifier (AR), and low threshold transient Ca2+ with maximal conductance of 60.5, 90, 5, 0.5, 0.005, and 0.5 mS/cm2 respectively. Leak current was set at 0.1 mS/cm^2^ for soma compartments, and 0.035 mS/cm^2^ for dendrites, except when altering input resistance where leak conductance was scaled by 0.75 to 1.45 times, corresponding to ±25% change in input resistance. Dendritic compartments had no Na+ conductance and had lower Ca2+ conductance of 0.15 mS/cm^2^. We removed the sodium current from dendrites, as TRN dendrites do not spike in recordings (Connelly et al., 2017), additionally the case of static dendrites were simpler and could be generalized to coupled neurons in other brain regions. Reversal potentials were 50 mV for sodium, -100 mV for potassium, 125 mV for calcium, -40 mV for AR and -75 mV for leak. Electrical synapses were modeled as a static conductance of the voltage difference between the coupled compartments of the TRN cells. Excitatory synapses were AMPAergic with reversal potential of 0 mV with rise and fall time kinetics of 5 ms and 35 ms respectively.

### Analysis

Coupling coefficients were measured by injecting hyperpolarizing current into the soma of one cell (A) and measuring the resulting current deflection in the soma of the other cell (B) compared to baseline (cc_AB_ = dV_B_/dV_A_), matching experimental methodology. We used 500 ms long square current injection and measured coupling in both directions between the cell pairs. The steady-state voltage during hyperpolarization was taken as the average voltage during the last 200 ms of stimulation. Coupling coefficient ratio was calculated as cc_12_/cc_21_.

To analyze latency modulation produced by electrical synapses we applied burst like EPSCs to distal dendrites of the model TRN cells and measured the time between onset of the first EPSC to the first action potential. Bursts consisted of 13 EPSCs of 1 µA amplitude with 5 ms ISI, 0.5 µA depolarizing current was applied to raise excitability of the cell model. Latency modulation was expressed as the difference in the change of latency between the two cells, compared to the latency of an uncoupled model cell.

## REFERENCES

Apostolides, P. F., & Trussell, L. O. (2013). Regulation of interneuron excitability by gap junction coupling with principal cells. Nature Neuroscience, 16(12), 1764–1772. https://doi.org/10.1038/nn.3569

Bukauskas, F. F., Elfgang, C., Willecke, K., & Weingart, R. (1995). Heterotypic gap junction channels (connexin26—Connexin32) violate the paradigm of unitary conductance. Pflügers Archiv European Journal of Physiology, 429(6), 870–872. https://doi.org/10.1007/BF00374812

Chagnaud, B. P., Perelmuter, J. T., Forlano, P. M., & Bass, A. H. (2021). Gap junction-mediated glycinergic inhibition ensures precise temporal patterning in vocal behavior. ELife, 10, e59390. https://doi.org/10.7554/eLife.59390

Choi, U., Wang, H., Hu, M., Kim, S., & Sieburth, D. (2021). Presynaptic coupling by electrical synapses coordinates a rhythmic behavior by synchronizing the activities of a neuron pair. Proceedings of the National Academy of Sciences, 118(20), e2022599118. https://doi.org/10.1073/pnas.2022599118

Chow, C. C., & Kopell, N. (2000). Dynamics of Spiking Neurons with Electrical Coupling. Neural Computation, 12(7), 1643–1678. https://doi.org/10.1162/089976600300015295

Connelly, W. M., Crunelli, V., & Errington, A. C. (2017). Variable Action Potential Backpropagation during Tonic Firing and Low-Threshold Spike Bursts in Thalamocortical But Not Thalamic Reticular Nucleus Neurons. The Journal of Neuroscience, 37(21), 5319–5333. https://doi.org/10.1523/JNEUROSCI.0015-17.2017

Curti, S., Hoge, G., Nagy, J. I., & Pereda, A. E. (2012). Synergy between Electrical Coupling and Membrane Properties Promotes Strong Synchronization of Neurons of the Mesencephalic Trigeminal Nucleus. Journal of Neuroscience, 32(13), 4341–4359. https://doi.org/10.1523/JNEUROSCI.6216-11.2012

Devor, A., & Yarom, Y. (2002). Electrotonic Coupling in the Inferior Olivary Nucleus Revealed by Simultaneous Double Patch Recordings. Journal of Neurophysiology, 87(6), 3048–3058. https://doi.org/10.1152/jn.2002.87.6.3048

Dugué, G. P., Brunel, N., Hakim, V., Schwartz, E., Chat, M., Lévesque, M., Courtemanche, R., Léna, C., & Dieudonné, S. (2009). Electrical Coupling Mediates Tunable Low-Frequency Oscillations and Resonance in the Cerebellar Golgi Cell Network. Neuron, 61(1), 126– 139. https://doi.org/10.1016/j.neuron.2008.11.028

Fortier, P. A. (2010). Detecting and estimating rectification of gap junction conductance based on simulations of dual-cell recordings from a pair and a network of coupled cells. Journal of Theoretical Biology, 265(2), 104–114. https://doi.org/10.1016/j.jtbi.2010.03.048

Fricker, B., Heckman, E., Cunningham, P. C., Wang, H., & Haas, J. S. (2021). Activity-dependent long-term potentiation of electrical synapses in the mammalian thalamus. Journal of Neurophysiology, 125(2), 476–488. https://doi.org/10.1152/jn.00471.2020

Fukuda, T. (2017). Structural organization of the dendritic reticulum linked by gap junctions in layer 4 of the visual cortex. Neuroscience, 340, 76–90. https://doi.org/10.1016/j.neuroscience.2016.10.050

Furshpan, E. J., & Potter, D. D. (1959). Transmission at the giant motor synapses of the crayfish. The Journal of Physiology, 145(2), 289–325. https://doi.org/10.1113/jphysiol.1959.sp006143

Galarreta, M., & Hestrin, S. (2002). Electrical and chemical synapses among parvalbumin fast-spiking GABAergic interneurons in adult mouse neocortex. Proceedings of the National Academy of Sciences, 99(19), 12438–12443. https://doi.org/10.1073/pnas.192159599

Gentet, L. J., & Ulrich, D. (2003). Strong, reliable and precise synaptic connections between thalamic relay cells and neurones of the nucleus reticularis in juvenile rats. The Journal of Physiology, 546(3), 801–811. https://doi.org/10.1113/jphysiol.2002.032730

Gutierrez, G. J., & Marder, E. (2013). Rectifying Electrical Synapses Can Affect the Influence of Synaptic Modulation on Output Pattern Robustness. Journal of Neuroscience, 33(32), 13238–13248. https://doi.org/10.1523/JNEUROSCI.0937-13.2013

Gutierrez, G. J., O’Leary, T., & Marder, E. (2013). Multiple Mechanisms Switch an Electrically Coupled, Synaptically Inhibited Neuron between Competing Rhythmic Oscillators. Neuron, 77(5), 845–858. https://doi.org/10.1016/j.neuron.2013.01.016

Haas, J. S. (2015). A new measure for the strength of electrical synapses. Frontiers in Cellular Neuroscience, 9. https://doi.org/10.3389/fncel.2015.00378

Haas, J. S., Zavala, B., & Landisman, C. E. (2011). Activity-Dependent Long-Term Depression of Electrical Synapses. Science, 334(6054), 389–393. https://doi.org/10.1126/science.1207502

Hoehne, A., McFadden, M. H., & DiGregorio, D. A. (2020). Feed-forward recruitment of electrical synapses enhances synchronous spiking in the mouse cerebellar cortex. ELife, 9, e57344. https://doi.org/10.7554/eLife.57344

Hoge, G. J., Davidson, K. G. V., Yasumura, T., Castillo, P. E., Rash, J. E., & Pereda, A. E. (2011). The extent and strength of electrical coupling between inferior olivary neurons is heterogeneous. Journal of Neurophysiology, 105(3), 1089–1101. https://doi.org/10.1152/jn.00789.2010

Johnson, B. R., Peck, J. H., & Harris-Warrick, R. M. (1993). Amine modulation of electrical coupling in the pyloric network of the lobster stomatogastric ganglion. Journal of Comparative Physiology A, 172(6). https://doi.org/10.1007/BF00195397

Lang, E. J., Sugihara, I., & Llinas, R. (1996). GABAergic Modulation of Complex Spike Activity by the Cerebellar Nucleoolivary Pathway in Rat. Journal of Neurophysiology, 76(1), 21.

Lee, S.-C., Patrick, S. L., Richardson, K. A., & Connors, B. W. (2014). Two Functionally Distinct Networks of Gap Junction-Coupled Inhibitory Neurons in the Thalamic Reticular Nucleus. Journal of Neuroscience, 34(39), 13170–13182. https://doi.org/10.1523/JNEUROSCI.0562-14.2014

Lewis, T. J., & Rinzel, J. (2003). Dynamics of Spiking Neurons Connected by Both Inhibitory and Electrical Coupling. Journal of Computational Neuroscience, 14, 283–309.

Li, X., Olson, C., Lu, S., Kamasawa, N., Yasumura, T., Rash, J. E., & Nagy, J. I. (2004). Neuronal connexin36 association with zonula occludens-1 protein (ZO-1) in mouse brain and interaction with the first PDZ domain of ZO-1. European Journal of Neuroscience, 19(8), 2132–2146. https://doi.org/10.1111/j.0953-816X.2004.03283.x

Liu, P., Chen, B., Mailler, R., & Wang, Z.-W. (2017). Antidromic-rectifying gap junctions amplify chemical transmission at functionally mixed electrical-chemical synapses. Nature Communications, 8(1), 14818. https://doi.org/10.1038/ncomms14818

Llinas, R., Baker, R., & Sotelo, C. (1974). Electrotonic coupling between neurons in cat inferior olive. Journal of Neurophysiology, 37(3), 560–571. https://doi.org/10.1152/jn.1974.37.3.560

Long, M. A. (2004). Small Clusters of Electrically Coupled Neurons Generate Synchronous Rhythms in the Thalamic Reticular Nucleus. Journal of Neuroscience, 24(2), 341–349. https://doi.org/10.1523/JNEUROSCI.3358-03.2004

Mainen, Z. F., & Sejnowski, T. J. (1996). Influence of dendritic structure on firing pattern in model neocortical neurons. Nature, 382(6589), 363–366. https://doi.org/10.1038/382363a0

Mann-Metzer, P., & Yarom, Y. (1999). Electrotonic Coupling Interacts with Intrinsic Properties to Generate Synchronized Activity in Cerebellar Networks of Inhibitory Interneurons. The Journal of Neuroscience, 19(9), 3298–3306. https://doi.org/10.1523/JNEUROSCI.19-09-03298.1999

Marsh, A. J., Michel, J. C., Adke, A. P., Heckman, E. L., & Miller, A. C. (2017). Asymmetry of an Intracellular Scaffold at Vertebrate Electrical Synapses. Current Biology, 27(22), 3561-3567.e4. https://doi.org/10.1016/j.cub.2017.10.011

McCormick, D. A., & Huguenard, J. R. (1992). A model of the electrophysiological properties of thalamocortical relay neurons. Journal of Neurophysiology, 68(4), 1384–1400. https://doi.org/10.1152/jn.1992.68.4.1384

Palacios-Prado, N., Hoge, G., Marandykina, A., Rimkute, L., Chapuis, S., Paulauskas, N., Skeberdis, V. A., O’Brien, J., Pereda, A. E., Bennett, M. V. L., & Bukauskas, F. F. (2013). Intracellular Magnesium-Dependent Modulation of Gap Junction Channels Formed by Neuronal Connexin36. Journal of Neuroscience, 33(11), 4741–4753. https://doi.org/10.1523/JNEUROSCI.2825-12.2013

Palumbos, S., Skelton, R., McWhirter, R., Mitchell, A., Swann, I., Heifner, S., Von Stetina, S., & Miller, D. M. (2021). CAMP controls a trafficking mechanism that directs the neuron specificity and subcellular placement of electrical synapses [Preprint]. Neuroscience. https://doi.org/10.1101/2021.05.12.443836

Pfeuty, B., Mato, G., Golomb, D., & Hansel, D. (2005). The Combined Effects of Inhibitory and Electrical Synapses in Synchrony. Neural Computation, 17(3), 633–670. https://doi.org/10.1162/0899766053019917

Pham, T., & Haas, J. S. (2018). Electrical synapses between inhibitory neurons shape the responses of principal neurons to transient inputs in the thalamus: A modeling study. Scientific Reports, 8(1), 7763. https://doi.org/10.1038/s41598-018-25956-x

Pham, T., & Haas, J. S. (2019). Electrical synapses regulate both subthreshold integration and population activity of principal cells in response to transient inputs within canonical feedforward circuits. PLOS Computational Biology, 15(2), e1006440. https://doi.org/10.1371/journal.pcbi.1006440

Phelan, P., Goulding, L. A., Tam, J. L. Y., Allen, M. J., Dawber, R. J., Davies, J. A., & Bacon, J. P. (2008). Molecular Mechanism of Rectification at Identified Electrical Synapses in the Drosophila Giant Fiber System. Current Biology, 18(24), 1955–1960. https://doi.org/10.1016/j.cub.2008.10.067

Rash, J. E., Curti, S., Vanderpool, K. G., Kamasawa, N., Nannapaneni, S., Palacios-Prado, N., Flores, C. E., Yasumura, T., O’Brien, J., Lynn, B. D., Bukauskas, F. F., Nagy, J. I., & Pereda, A. E. (2013). Molecular and Functional Asymmetry at a Vertebrate Electrical Synapse. Neuron, 79(5), 957–969. https://doi.org/10.1016/j.neuron.2013.06.037

Sevetson, J., & Haas, J. S. (2015). Asymmetry and modulation of spike timing in electrically coupled neurons. Journal of Neurophysiology, 113(6), 1743–1751. https://doi.org/10.1152/jn.00843.2014

Shui, Y., Liu, P., Zhan, H., Chen, B., & Wang, Z.-W. (2020). Molecular basis of junctional current rectification at an electrical synapse. Science Advances, 6(27), eabb3076. https://doi.org/10.1126/sciadv.abb3076

Snipas, M., Rimkute, L., Kraujalis, T., Maciunas, K., & Bukauskas, F. F. (2017). Functional asymmetry and plasticity of electrical synapses interconnecting neurons through a 36-state model of gap junction channel gating. PLOS Computational Biology, 13(4), e1005464. https://doi.org/10.1371/journal.pcbi.1005464

Srinivas, M., Rozental, R., Kojima, T., Dermietzel, R., Mehler, M., Condorelli, D. F., Kessler, J. A., & Spray, D. C. (1999). Functional Properties of Channels Formed by the Neuronal Gap Junction Protein Connexin36. The Journal of Neuroscience, 19(22), 9848–9855. https://doi.org/10.1523/JNEUROSCI.19-22-09848.1999

Szoboszlay, M., Lőrincz, A., Lanore, F., Vervaeke, K., Silver, R. A., & Nusser, Z. (2016). Functional Properties of Dendritic Gap Junctions in Cerebellar Golgi Cells. Neuron, 90(5), 1043–1056. https://doi.org/10.1016/j.neuron.2016.03.029

Teubner, B., Degen, J., Söhl, G., Güldenagel, M., Bukauskas, F. F., Trexler, E. B., & Willecke, K. (2013). Functional Expression of the Murine Connexin 36 Gene Coding for a Neuron-Specific Gap Junctional Protein. 21.

Traub, R. D., Contreras, D., Cunningham, M. O., Murray, H., LeBeau, F. E. N., Roopun, A., Bibbig, A., Wilent, W. B., Higley, M. J., & Whittington, M. A. (2005). Single-Column Thalamocortical Network Model Exhibiting Gamma Oscillations, Sleep Spindles, and Epileptogenic Bursts. Journal of Neurophysiology, 93(4), 2194–2232. https://doi.org/10.1152/jn.00983.2004

Veruki, M. L., & Hartveit, E. (2002). AII (Rod) Amacrine Cells Form a Network of Electrically Coupled Interneurons in the Mammalian Retina. Neuron, 33(6), 935–946. https://doi.org/10.1016/S0896-6273(02)00609-8

Vervaeke, K., Lőrincz, A., Gleeson, P., Farinella, M., Nusser, Z., & Silver, R. A. (2010). Rapid Desynchronization of an Electrically Coupled Interneuron Network with Sparse Excitatory Synaptic Input. Neuron, 67(3), 435–451. https://doi.org/10.1016/j.neuron.2010.06.028

Zolnik, T. A., & Connors, B. W. (2016). Electrical synapses and the development of inhibitory circuits in the thalamus: Electrical synapses and thalamic development. The Journal of Physiology, 594(10), 2579–2592. https://doi.org/10.1113/JP271880

